# ProtNHF: Neural Hamiltonian Flows for Controllable Protein Sequence Generation

**DOI:** 10.64898/2026.03.04.709305

**Authors:** Bharath Raghavan, David M. Rogers

**Affiliations:** National Center for Computational Sciences, Oak Ridge National Laboratory, Oak Ridge, United States of America

## Abstract

Controllable protein sequence generation remains a central challenge in computational protein design, as most existing approaches rely on retraining, classifier guidance, or architectural modification to impose conditioning. Here we introduce ProtNHF, a generative model that enables continuous, quantitative control over sequence-level properties through analytical bias functions applied exclusively at inference time. ProtNHF builds on neural Hamiltonian flows, where a lightweight Transformer-based potential energy function, inspired by ESM-2, is combined with an explicit kinetic term to define Hamiltonian dynamics in a continuous relaxation of protein sequence space. The model learns a symplectic transport map from a latent Gaussian distribution to protein sequence embeddings via deterministic leapfrog integration, enabling efficient and expressive sampling. In the unconditional setting, generated sequences achieve competitive quality as measured by ESM-2 pseudo-perplexity and AlphaFold2 pLDDT confidence scores. A key advantage of the Hamiltonian formulation is its additive energy structure, which permits external bias potentials to be incorporated directly into the Hamiltonian at inference time without modifying or retraining the learned model. This casts controllable generation in a classical molecular modeling paradigm, where desired properties are enforced by explicit energy shaping. We demonstrate smooth, predictable, and approximately monotonic control over amino acid composition and global properties such as net charge by introducing simple analytical bias terms, including residue-specific chemical potentials and harmonic constraints. The bias strength modulates the values of these properties in generated sequences continuously while preserving structural plausibility and diversity. ProtNHF thus provides a flexible base distribution that can be steered toward different compositional regimes using transparent, physically interpretable energy terms, establishing a general framework for inference-time programmable protein sequence generation.

## 1 Introduction

Predicting protein sequences that reliably fold into valid three-dimensional structures remains a fundamentally challenging problem.[1, 2, 3] Recent advances in neural network architectures and modern generative modeling paradigms such as diffusion-based, autoregressive, and flow-based approaches have substantially expanded our ability to model complex data distributions.[4, 5, 6, 7, 8] These developments have enabled the generation of diverse and structurally plausible proteins, with applications ranging from enzyme engineering to therapeutic design.[9, 10, 11, 12, 13, 14, 15]

While unconditional protein generation already has significant practical value, many real-world protein engineering tasks require a more refined capability: the controlled generation of sequences that satisfy specified biochemical or compositional constraints. This constitutes protein engineering and this need is particularly acute in the design of artificial proteins and functional biomolecules.[16, 17, 18] Here global sequence-level properties such as amino acid composition, net charge, solubility, hydrophobicity, or disorder propensity must often be tuned to meet experimental or biophysical requirements.[19, 20, 21, 22]

Most existing approaches to controllable protein sequence generation rely on explicit conditioning mechanisms (e.g., taxonomic/structural control tags) or objective-specific fine-tuning, either by modifying model architecture or by leveraging auxiliary predictors during training or inference[23, 24, 25, 26, 27, 28] Although effective, these approaches typically require retraining or architectural modification for each new target property, limiting flexibility and increasing computational cost. Structure-first generative models such as Chroma[29] introduce powerful programmable capabilities in structure space, but operate via diffusion over three-dimensional conformations and do not directly provide fine-grained, continuous control over global sequence-level statistics. As a result, achieving predictable and quantitative modulation of compositional properties without retraining remains an open challenge.

The recently introduced generative models of neural Hamiltonian flows or NHFs hold great promise in this regard. First introduced in the works of Higgins *et. al*.[30] and Toth *et. al*.[31], NHFs are a type of normalizing flow which transforms an initial Gaussian distribution in phase space into the final distribution using a sequence of invertible, volume-preserving transformations derived from Hamiltonian dynamics. Recent works have empirically demonstrated their ability to model complex distributions involving applications in image generation, cosmology and solving the 1D Vlasov-Poisson Equations. [32, 33] Related Hamiltonian-based generative formulations, such as Hamiltonian Score Matching, have also recently been proposed, further underscoring the broader applicability of Hamiltonian dynamics in generative modeling.[34] However, no study has applied NHFs to more complex generative problems like protein sequence generation.

In this work, we introduce **ProtNHF**, a generative model for protein sequences that enables continuous, quantitative control over sequence-level properties via analytical bias functions applied at inference time. ProtNHF builds upon the framework of NHFs, where the learned potential energy function in the NHF is parameterized by a lightweight Transformer architecture[35] inspired from the ESM-2 model[11] and combined with an explicit kinetic energy term to define Hamiltonian dynamics in a continuous relaxation of protein sequence space.

A key advantage of the Hamiltonian formulation is its additive energy structure.[36] This means that external bias potentials can be incorporated directly into the Hamiltonian at inference time without modifying or retraining the learned model. This allows controllable generation to be formulated within a classical molecular modeling paradigm[37, 38], where desired properties are enforced by explicitly modifying the energy landscape.

Our central contribution is to demonstrate that (i) NHFs can be applied to model complex data distributions, like the protein sequence space; and (ii) this formulation enables smooth, predictable, and approximately monotonic control over amino acid composition and global sequence-level properties. By introducing simple analytical bias terms, such as residue-specific chemical potentials inspired by molecular simulations, we show that the fraction of desired residues and properties such as net charge can be increased or decreased continuously by tuning a single scalar bias parameter. This control operates globally across the sequence and preserves diversity without requiring retraining. Furthermore, secondary structure diversity can also be manipulated with these analytical biases. ProtNHF thus serves as a flexible base generative model that can be steered towards different compositional regimes using transparent, physically interpretable energy terms, providing a general framework for inference-time programmable protein sequence generation.

## 2 Background

In classical mechanics, dynamics over the phase space variables **z** ≡ (**q, p**) can be written as

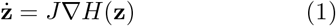

Here, 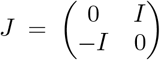 is the symplectic matrix and *H* is the Hamiltonian[36], defined as the scalar total energy of the system. The latter can be expressed as the sum of potential *V* (solely dependent on the generalized coordinates **q**) and kinetic energy *K* (solely dependent on the generalized momenta **p** and mass *m* as *K*(**p**) = **p**^**2**^/2*m*). The Hamiltonian dynamics in Equation 1 can then be written as

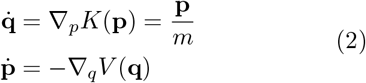

Neural Hamiltonian Flows or NHFs are a class of generative models mapping between samples from the complex target distribution (**q**_*T*_, **p**_*T*_) ∼ π_T_ onto a known latent distribution (**q**_0_, **p**_0_) ∼ π_0_ through a series of discretized Hamiltonian maps[30]

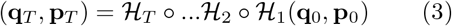

where each Hamiltonian map ℋ_*i*_ is the *i*^*th*^ step of the discretized form of the continuous Hamiltonian equations (Equation 2) using the symplectic Leapfrog integrator[39]. The discretization is done for *T* steps with timestep Δt. The potential energy is expressed as a a learned neural network *V*_*θ*_(**q**) and the kinetic energy is fixed as *K*(**p**) = **p**^2^*/*2*M*_*θ*_ in terms of a positive-definite, learned mass matrix *M*_*θ*_ as in ref [32].

In order to avoid having to artificially split the density into two disjoint sets **z** ≡ (**q, p**), the momentum variable **p** is treated as a latent variables in a similar interpretation to Hamiltonian Monte Carlo.[39] This involves an encoder made of two neural networks 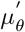 and 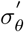 which are used to sample **p** as

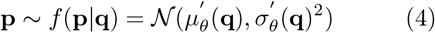

The training is performed in the ‘forward’ direction or by running the Hamiltonian transformation from (**q**_*T*_, **p**_*T*_) → (**q**_0_, **p**_0_).

The model is trained by maximizing the NHF loss function given as

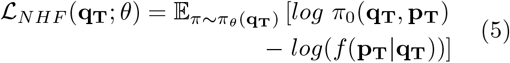

The latent distribution *π*_0_ can be fixed as any convenient distribution. In practice, a Gaussian distribution of fixed mean and standard deviation is used. The Hamiltonian mapping between (**q**_*T*_, **p**_*T*_) → (**q**_0_, **p**_0_) is invertible. This means that having learned the transformation, one can reverse the sign of the timestep Δt and use the same Hamiltonian to transform the latent distribution into the target distribution (**q**_0_, **p**_0_) → (**q**_T_, **p**_*T*_). Thus, new samples from *π*_*T*_ can be generated by starting with samples from the latent Gaussian distribution *π*_0_ and running the reverse or ‘sampling’ process.

## 3 Methods

In this section, we describe the changes introduced by us to the general NHF scheme to allow for protein sequence generation.

### 3.1 Designing the Embedding Space for Protein Sequences

A protein is represented as a sequence of amino acids *X* = [*a*_1_, *a*_2_, …, *a*_*L*_] ≡ [**x**_1_, **x**_2_, …, **x**_*L*_] of length *L*, where each element *a*_*i*_ is one of the 20 standard amino acids. Each *a*_*i*_ is represented by a one-hot encoding vector **x**_*i*_. In this work, we use NHFs for de novo protein sequence design by learning the data distribution over all possible protein sequences and to sample novel sequences *X* from this distribution. The flow acts over the phase space variables (**q, p**), which must exist on continuous space. However, a protein sequence is a permutation of 20 standard amino acid residues. These are categorical (hence, discrete) values, and an NHF cannot be applied on them directly. To solve this, we represent the coordinate variable for the *i*^*th*^ residue in the sequence **q**_*i*_ as the one-hot encoding representation of that residue **x**_*i*_ mapped onto a continuous space using the argmax flow technique.[40] This is done by first choosing random Gaussian noise *u* as

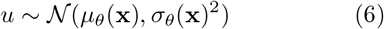

where *µ*_*θ*_ and *σ*_*θ*_ are neural networks. Then a threshold variable is written as 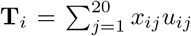 where, *i* = 1, …, *L* and the summation is performed over the 20 amino acid classes. Finally, the protein embeddings in continuous space **q** are defined as

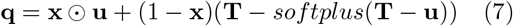

The mapping can be simply reversed by **x** = *argmax*(**q**). This is advantageous as it is a bijective mapping between protein sequences **x** and **q**, allowing it to be exactly reversible.

Looking at Equation 5, the new loss function for the complete flow becomes:

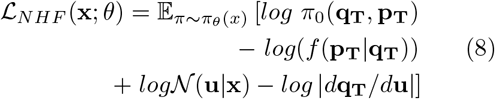

### 3.2 Conditional Generation

In this work, we argue that NHFs provide a powerful, yet interpretable way to generate conditional samples without additional training. Conditional sampling can be achieved by introducing an additional bias energy term into the Hamiltonian during the reverse process only, directly analogous to performing biased molecular dynamics simulations in the NVE ensemble.[37] For this, we introduce a smooth bias potential *U* (**q**) defined over the latent coordinate space. The potential encodes a design objective, such as penalizing specific residue embeddings or enforcing physicochemical constraints. Given the original Hamiltonian *H*(**q, p**), with the learned potential energy *V*_*θ*_(**q**), we define the biased Hamiltonian as *H*^*b*^(**q, p**) as

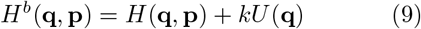

where *k* controls the conditioning or bias strength. Because the bias depends only on the coordinate variable **q**, it modifies only the force term in the momentum equation while preserving Hamiltonian structure. Biased Hamiltonian dynamics can be then be written in the symplectic matrix notation as

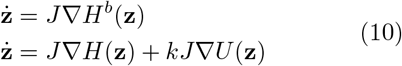

Thus, this energy-shaped conditioning can be interpreted as a smooth perturbation of the original, learned Hamiltonian system. For sufficiently small *k*, we can approximately write the relationship between biased Hamiltonian mapping 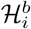 to the unbiased case ℋ_*i*_ as

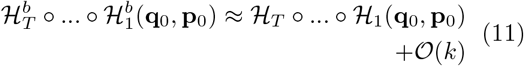

The biased Hamiltonian flows define a new generative model corresponding to a continuous deformation of the original dynamics; the strength of which is given by *k*. Importantly, the original potential energy *V*_*θ*_(**q**) learned during the training phase does not require any retraining for us to define this new conditional flow. While post-training guidance has been explored in diffusion and neural ODE frameworks, our approach leverages energy shaping within a symplectic Hamiltonian system. This ensures that conditioning preserves invertibility, phase-space volume, and geometric structure, yielding a controlled deformation of the learned flow.

Crucially, since the NHF is physically interpretable, we can use analytical functions to bias the simulations. Our ProtNHF model has been designed to support three forms of biasing potentials: Coulomb, Gaussian and Harmonic. *U* (**q**) can be represented as a Coulombic potential centered on **r** in coordinate space:

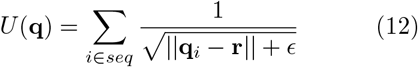

A value of *ϵ* = 10^™6^ is added here for numerical stability. This would steer the dynamics in the sampling stage away from the point **r**. *U* (**q**) can be represented as a Gaussian potential in a similar manner:

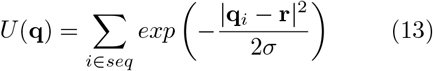

Here, *σ* is an additional parameter determining the width of the Gaussian potential.

If **r** is fixed as the embedding vector corresponding to a certain residue, these biasing functions can be used to generate sequences with more or less of the chosen residue.

The harmonic potential can also be used. It can be applied at a particular residue in the *n*^*th*^ position of the sequence:

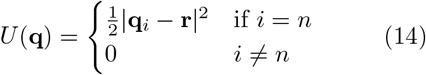

This would force the generative flow towards a protein with a chosen residue (the one-hot encoding of which is **q**_*c*_) in the *n*^*th*^ position. It can also be used to bias the generation towards proteins with certain global properties. For example, given a continuous function of the sequence embeddings *F* (**q**) that outputs a scalar, we can force the generation towards *F*^∗^ by adding the following bias:

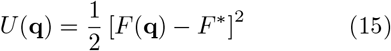

### 3.3 Model Architecture

The architecture of ProtNHF is depicted in Figure 1. The Hamiltonian equations of motion are discretized using the leapfrog integrator for 4 steps and a step size of 0.05. The encoders for Gaussian noise in Equation 6, *µ*_*θ*_ and *σ*_*θ*_, are represented as a single feed forward neural network with one 128-dimensional hidden layer and an output layer with two heads. Similarly for 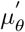 and 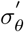 in Equation 4. For the latent distribution, a Gaussian with mean of 0 and standard deviation of 0.7 was found to exhibit the best generative performance, and was used in this work.

**Figure 1:**
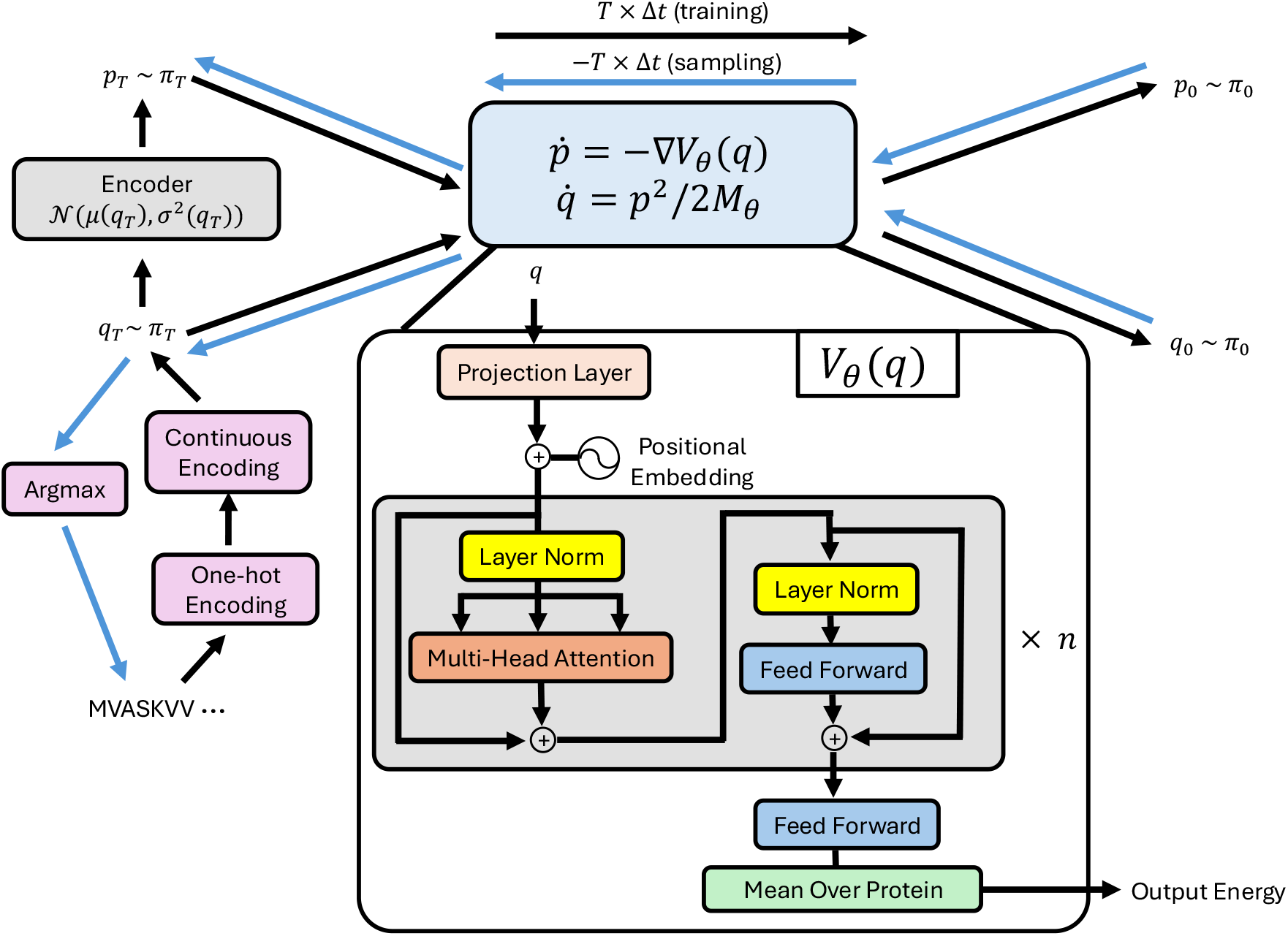
Overview of the ProtNHF model architecture. It consists of an NHF, discretized with the Leapfrog algorithm for *T* steps and a transformer *V*_*θ*_(**q**) with *n* heads used to calculate the ‘energy’ of the protein sequences.

An energy function is required to fully describe the Hamiltonian in the NHF. In order to capture the long-range interactions among the residues in a protein sequence, a multi-headed transformer is used. The number of parameters chosen for the transformer was taken to correspond to the 8M ESM-2 model (number of embedding dimensions as 320, hidden dimensions of feed forward layer as 1280, attention heads as 20 and hidden layers as 6).[11] The standard attention mechanism from Vaswani *et. al*.[35] would scale quadratically and would be prohibitively expensive to implement within a normalizing flow scheme. Hence, we opt to use the performer architecture, as a variant of the linear attention mechanism.[41]

The input to the transformer is the continuous protein embedding **q**. However, since each residue in the protein is only a 20-dimensional vector, the expressibility of the transformer is severely limited and cannot be directly used as input to the transformer. Therefore, a projection layer (which is just a single feed-forward layer) is used to project the vectors into a 320-dimensional space. Sinusoidal positional embeddings are then added to the projected outputs to allow the transformer to capture positional information. Prior to adding the positional information, the embeddings are scaled by multiplying them with the square root of the number of embedding dimensions before feeding them into the transformer.

The output of the transformer is also a 320-dimensional vector. These are passed to a feed-forward network to obtain a 1-dimensional vector for each residue, which are subsequently mean pooled over the protein sequences to get a value per protein. This is the ‘energy’ of the protein. The force is obtained by calculating the gradient of this energy with respect to **q** using the *torch*.*autograd*.*grad* feature of PyTorch.

## 4 Experiments and Results

Our ProtNHF model is trained with a learning rate of 10^-4^, warmed up over the first 5 epochs. A cosine scheduler is then used to decay the learning rate to 0 in 650 epochs. The dataset used consist of ∼90,000 sequences from the UniProtKB database, with proteins of lengths in the range of 10 to 128.[42] We assess the sequence quality of a protein sequences *X* = [*a*_1_, *a*_2_, …, *a*_*L*_] generated by the model, under both unconditional and conditional regimes, by three main criteria:

**ESM-2 pseudo-perplexity.** ESM-2 is a state-of-the-art protein language model that uses a BERT-style bidirectional transformer architecture to learn structural and evolutionary patterns directly from protein sequences. It was trained on millions of sequences from UniRef.[43] ESM-2 pseudoperplexity (ESM-2 pppl) measures how well a given protein sequence aligns with the patterns the model has learned from the training data. We calculate ESM-2 pppl with ESM-2 650M[11] by masking each amino acid of the protein sequence and predicting it considering all the other amino acids in the sequence, and calculating the value with the equation:

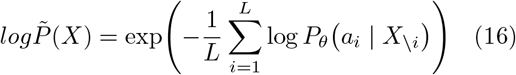

where *X*_\*i*_ denotes the sequence with residue *a*_*i*_ masked, and *P*_*θ*_ is the conditional distribution predicted by ESM-2.

Lower pseudo-perplexity values indicate that a sequence is more consistent with the distribution learned by the language model, and hence is more ‘protein-like’. A pppl value of 18 or above corresponds to a noisy or random sequence.

**Percentage low-complexity regions.** Low-complexity regions were identified using a sliding-window Shannon entropy criterion. The en-tropy was computed for each window *W*_*i*_ = (*a*_*i*_, …, *a*_*i*+*w*™1_) of size *w* = 12 as *H*(*W*_*i*_).

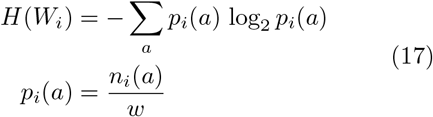

Windows with *H*(*W*_*i*_) < *H*_*th*_ = 2.2 *bits* were classified as low-complexity, and all residues covered by such windows were labeled accordingly. The percentage of low-complexity regions in the protein sequence (% LCRs) was defined as the fraction of residues labeled as low-complexity, reported as a percentage.

**Alphafold predicted local Distance Difference.** Structural confidence of predicted protein structures was evaluated using predicted Local Distance Difference Test (pLDDT) scores produced by AlphaFold.[44] For each residue, AlphaFold predicts a pLDDT score ranging from 0 to 100, which estimates the expected local structural accuracy of that residue. Higher pLDDT values correspond to higher confidence in the predicted local structure. The pLDDT for a protein sequence is calculated by taking the mean of the pLDDT values for all α-carbons. pLDDT scores above approximately 70 are generally associated with well-structured regions, whereas pLDDT values below 50 often indicate intrinsically disordered or flexible regions. Mean pLDDT is commonly used as a scalar proxy for foldability and structural confidence in computational protein design and sequence generation studies.

### 4.1 Unconditional Generation

The mean ESM-2 pppl and AlphaFold2 pLDDT for proteins of various lengths generated by ProtNHF model is plotted in Figure 2a. State-of-the-art models predict sequences with an average ESM-2 pppl in the range of 11–12.[12, 24, 13] The predictions for sequence length 20 from our model approach this range; however, the pppl values quickly diverge as sequence length increases. By length 50, pppl values are close to noise levels and do not correspond to clearly valid protein sequences. This trend is expected given that we used a relatively small transformer, equivalent to the smallest ESM-2 model, as our potential energy function and trained on a modest dataset (∼90,000 sequences). It can be anticipated that larger, more expressive transformers capable of capturing long-range dependencies would improve absolute performance. Despite this limitation, the model’s performance at shorter lengths exceeds initial expectations, indicating that NHFs can capture meaningful structure in the protein sequence landscape.

**Figure 2:**
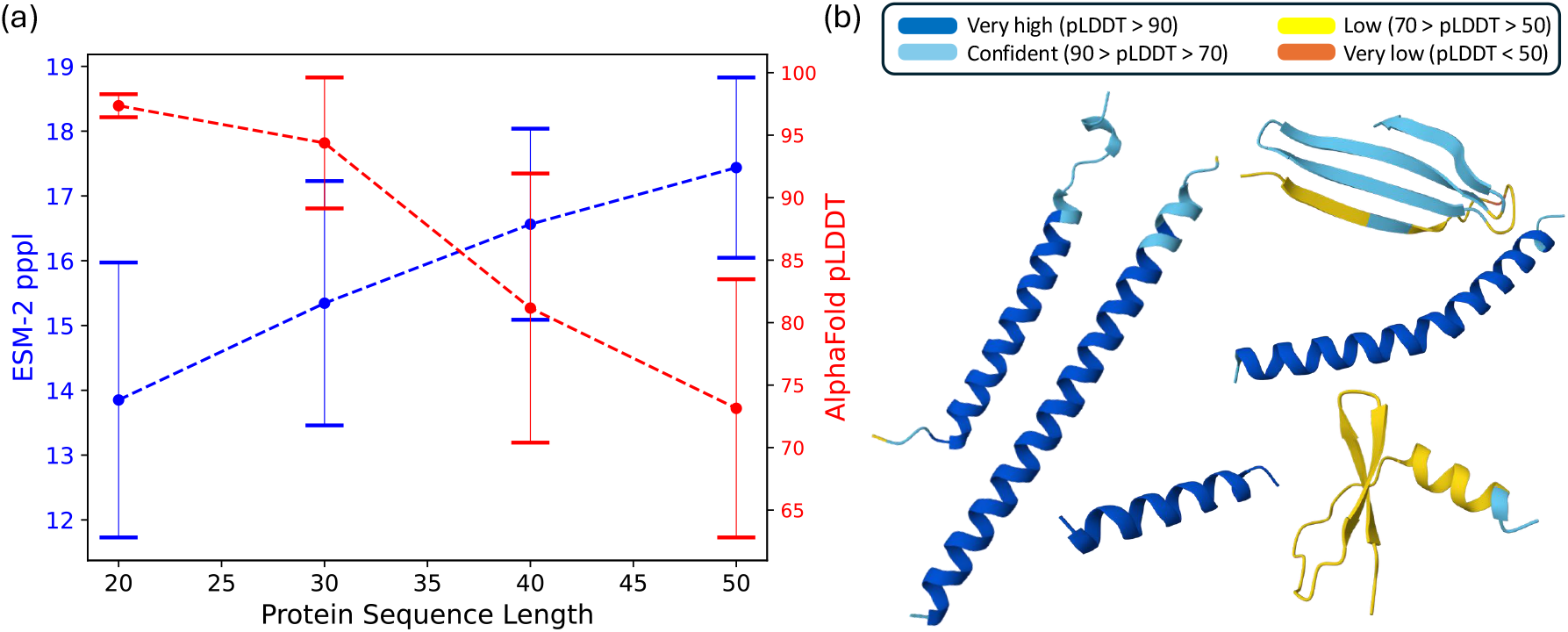
(a) Mean ESM-2 pppl and AlphaFold pLDDT for 500 and 30 protein sequences generated at various lengths by our NHF. Error bars represent one standard deviation across the independently generated sequences per length. (b) Representative folded structure by AlphaFold for sequences generated by ProtNHF.

In contrast to pppl, AlphaFold2 structural confidence remains relatively high for longer sequences. For lengths 20–30, the mean pLDDT lies between 90–100, indicating that the majority of generated sequences fold into highly confident structures. At length 40, the mean pLDDT decreases to approximately 80, and at length 50 it remains around 75. While this decline parallels the increase in pppl, the predicted structures at longer lengths still exhibit moderate-to-high confidence. Representative folded structures are shown in Figure 2b. Most generated proteins adopt predominantly α-helical conformations. A smaller subset forms β-sheets, with pLDDT values in the range of approximately 50–85.

Even in these cases, secondary structure elements are well defined, though global packing confidence decreases. These results suggest that although sequence likelihood deteriorates with length, the model continues to generate structurally coherent proteins up to lengths of 40-50 residues.

To further characterize sequence realism beyond language-model likelihood and structural confidence, we analyzed the prevalence of low-complexity regions (LCRs) in generated sequences. Excessive low-complexity content is a common degenerative mode in generative models, often reflecting repetitive or compositionally collapsed outputs. We computed the percentage of LCR residues for generated proteins of various lengths and examined the relationship between ESM-2 pppl and LCR fraction in Figure 3. For length-20 sequences, two distinct regimes are observed: a dominant cluster near zero LCR content and a secondary cluster near 100% low complexity. The latter corresponds to highly repetitive sequences, which are more likely at short lengths where combinatorial diversity is limited. As sequence length increases to 30, 40, and 50, the high-LCR cluster progressively diminishes and becomes negligible. Generated sequences instead concentrate near low LCR fractions even as pppl increases with length. The reduction of high-LCR sequences at longer lengths indicates that the model does not achieve favorable perplexity at short lengths just through trivial repetition.

**Figure 3:**
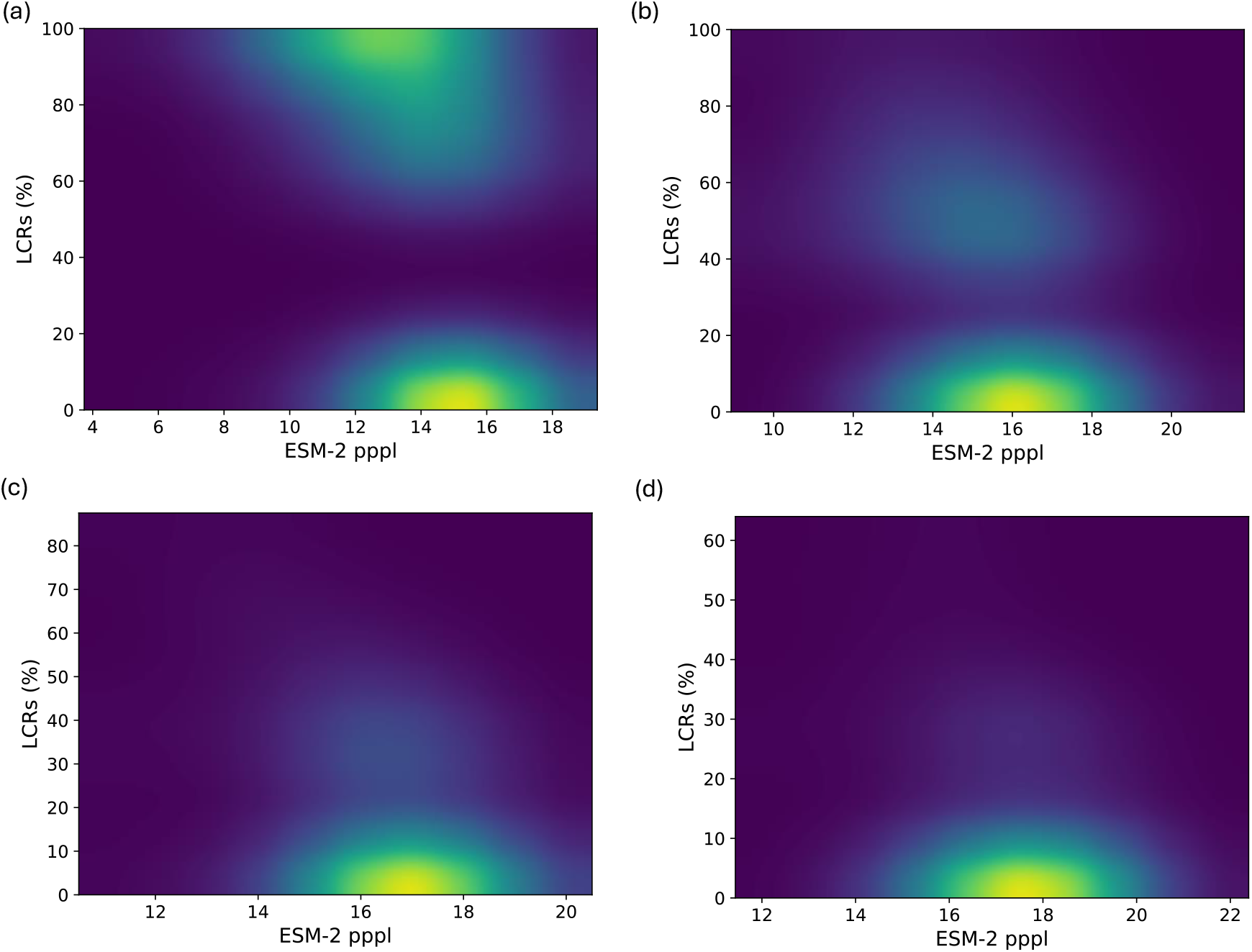
Histogram of ESM-2 pppl and % LCRs for 500 protein sequences generated by our NHF of lengths (a) 20, (b) 30, (c) 40 and (d) 50.

Taking all these results together suggests that while generative difficulty scales with length under the current model capacity, the learned Hamiltonian dynamics capture meaningful sequence statistics and produce structurally coherent outputs, particularly in the short length regime.

### 4.2 Conditional Generation using Analytical Biasing Functions

The most promising feature of NHFs is their ability to generate conditional outputs using analytical and physically interpretable biasing functions added during inference time, without the need for retraining. We demonstrate this for ProtNHF with the following experiments discussed below.

#### 4.2.1 Residue composition control

In the first experiment, protein sequences with reduced occurrences of Lys are generated. This is achieved by adding a Coulombic bias term, of the form shown in Equation 12, with **r** fixed as the one-hot encoding for Lys. Figure 4a shows the percentage reduction in the number of Lys and the ESM-2 pppl of length-20 protein sequences generated as a function of the strength of the Coulombic bias *k*. The number of Lys are a direct function of *k*, with only a slight increase in the ESM-2 pppl as *k* increases.

**Figure 4:**
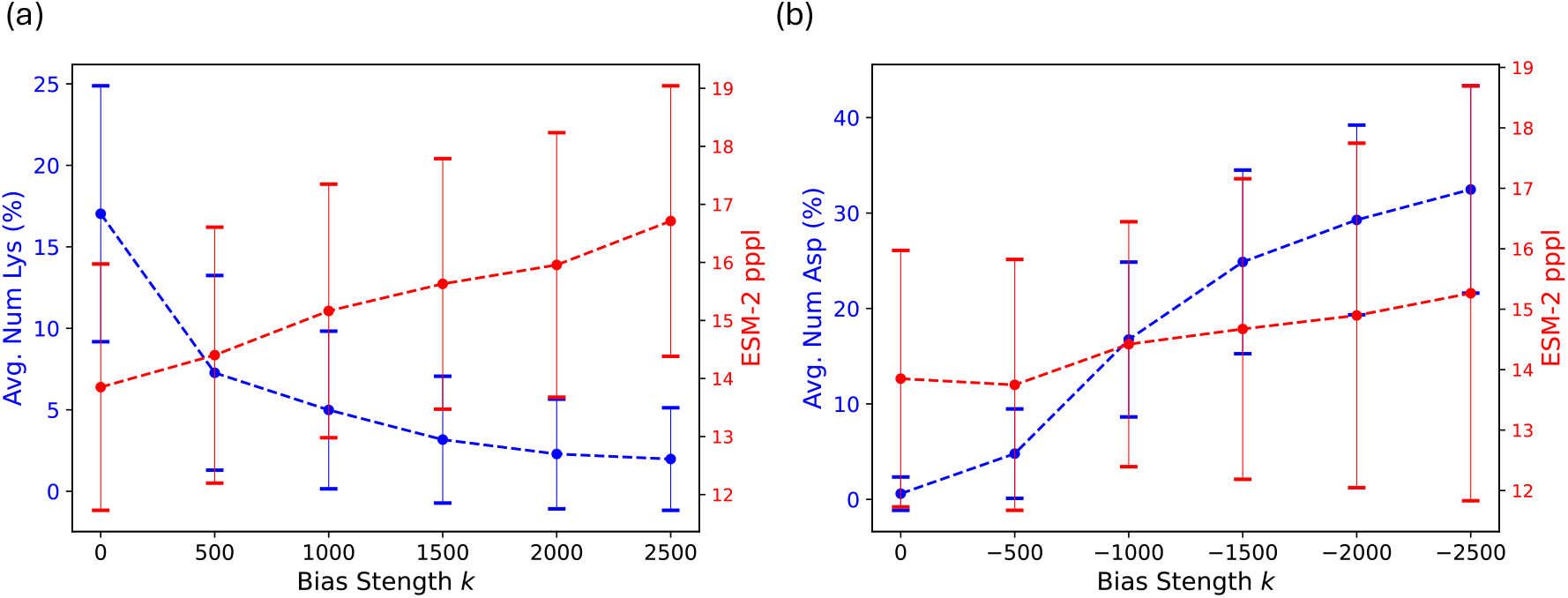
(a) Mean percentage reduction in Lys content and corresponding mean ESM-2 pppl for 500 generated length-20 protein sequences as a function of the Coulombic bias strength *k*. (b) Mean percentage increase in Asp content and corresponding mean ESM-2 pppl as a function of the Gaussian bias strength *k*. A more negative value implies a stronger attractive potential. Error bars represent one standard deviation across the independently generated sequences per *k* value.

In the second experiment, protein sequences with increased occurrences of Asp residues are generated. This is achieved by adding a Gaussian bias term of the form shown in Equation 13, with a negative bias strength, unity width and **r** fixed as the one-hot encoding for the residue Asp. Figure 4b shows the percentage increase in the number of Asp residues and change in the ESM-2 pppl of length-20 protein sequences generated as a function of the strength of the Gaussian bias *k*. Again, a more negative value of *k* leads to proteins with more aspartates, while still maintaining approximately the same value of ESM-2 pppl.

Sequences from selected bias strengths were folded in AlphaFold for both experiments (see cases A and B in Figure 6b). Comparing this with the AlphaFold pLDDT obtained for the unbiased case of length 20 (see Figure 2a), it can be seen that the bias does not adversely affect the structural confidence of the sequences. Thus, these two experiments show that NHFs allow continuous, quantitative control of protein residue composition at inference time through additive analytical energy terms, without retraining.

#### 4.2.2 Positional and global property control

Here we to demonstrate the use of the harmonic biasing function in ProtNHF. In molecular dynamics simulations, harmonic restraints are placed to fix the positions of certain atoms. In a direct parallel to this, it is possible to restraint ProtNHF to only generate proteins with certain residues in certain positions. As a third experiment, we demonstrate the usefulness of the harmonic restraint by forcing the model to generate protein sequences starting with the residue Met only. The restraint improves sequence plausibility and ESM-2 pppl scores by enforcing a biologically common N-terminal residue.

This is done by adding the bias as per Equation 14, with **r** fixed as the one-hot encoding of Met. Figure 5a shows the reduction in ESM-2 pppl values of the generated protein sequences. Furthermore, the % LCRs for each length is largely in line with the unbiased case. We also folded selected sequences (Figure 5b) and found that they produce greater diversity in secondary structure than the unbiased case. The mean AlphaFold pLDDT was also found to be higher than for unbiased sequences of length 40 (compare case C in Figure 6b with 2a). This indicates that residue control with ProtNHF can lead us to a pathway for generating protein sequences with greater structural and topological diversity.

**Figure 5:**
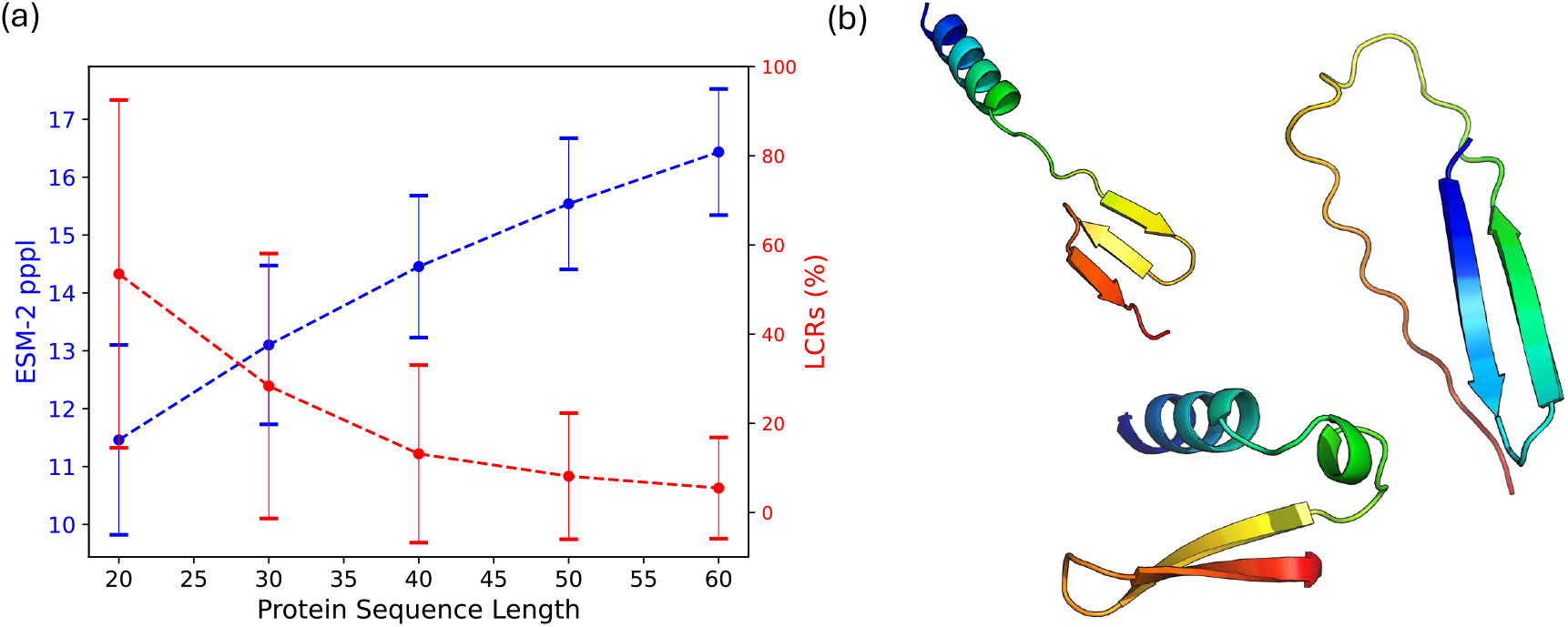
(a) Mean ESM-2 pppl and % LCRs for 500 length-20 protein sequences generated at various lengths by our NHF when applying a harmonic bias constraining sequences to begin with Met. Error bars represent one standard deviation across the independently generated sequences per length. (b) Representative folded structure by AlphaFold for sequences generated by constraining them to begin with Met.

Finally, we show how the bias as per Equation 15 can be used to generate protein sequences with various desired global properties. Here we take the example of total charge. This is done by setting *F* (**q**) in Equation 15 to a function that returns the total charge of the protein. To enable gradient-based control of sequence charge within the Hamiltonian flow, we compute a differentiable estimate of net charge directly from the sequence embeddings. For a sequence of length *L* with and embeddings **q**_*i,a*_ over amino acids *a* ∈ 1, …, 20, probabilities are obtained via a softmax

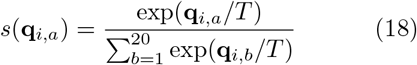

Each amino acid is assigned a fixed charge *c*(*a*) (here we use, +1 for Lys and Arg, −1 for Asp and Glu, 0.1 for His, and 0 for the rest). The net charge is then defined as the expectation under the softmax distribution as

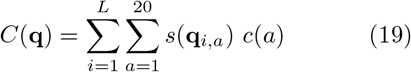

which is fully differentiable with respect to the sequence embeddings and can therefore be incorporated in the Hamiltonian. To test the effect of this bias, protein sequences of length 20 were generated with a target net charge of 0. Figure 6a depicts the mean net charge of the generated sequences moving closer to the target value as the bias strength *k* is increased, while the ESM-2 pppl remains largely constant. This shows that analytical biasing of the ProtNHF model during inference time does not only allow for direct residue control, but also generation of protein sequences with desired global properties. To further test this idea, we applied the net charge restraint at a target charge of −1 with a bias strength of *k* = 2000 and length 40. The resulting mean charge for 500 generates sequences was indeed close at −1.18 ± 0.58 with a mean ESM-2 pppl of 14.83 ± 2.02. In fact, the AlphaFold pLDDT was also found to be slightly higher than for unbiased sequences of length 40 (compare case D in Figure 6b with 2a), indicating greater structural stability of the generated sequences.

**Figure 6:**
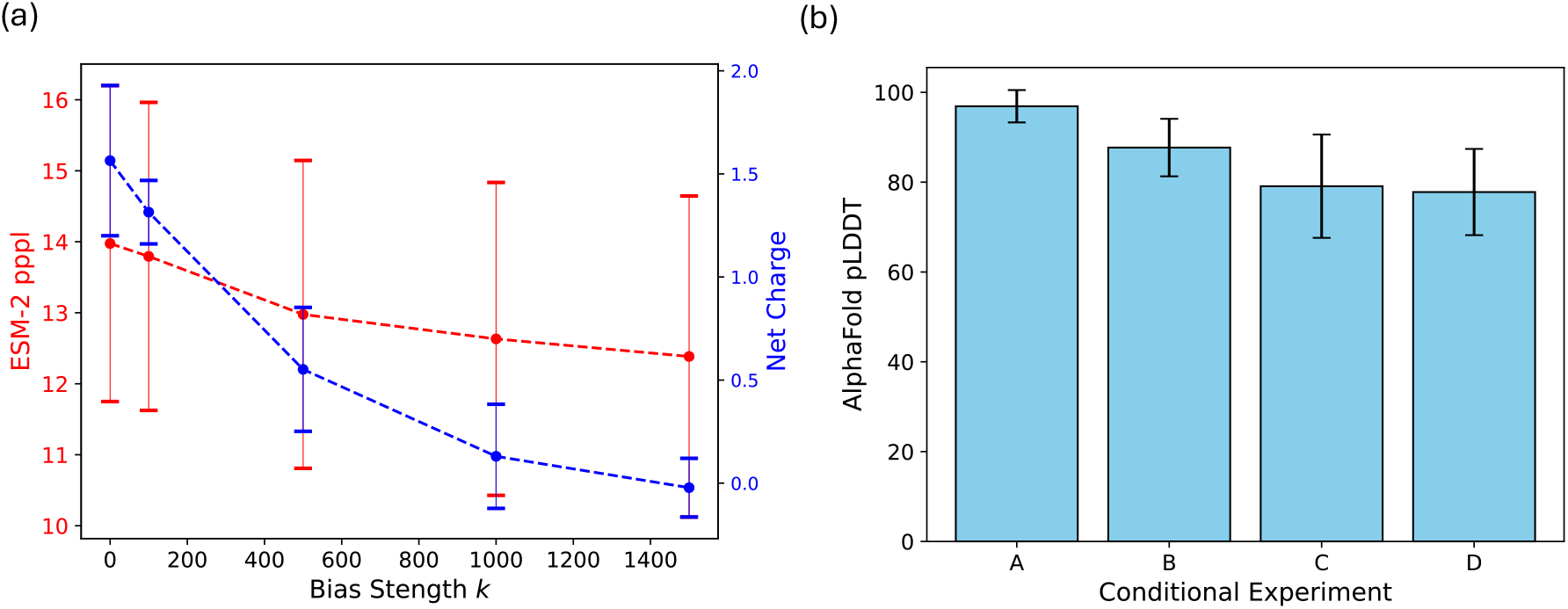
(a) Mean net charge of 500 generated length-20 protein sequences as a function of bias strength *k*, with a target net charge of zero. (b) Mean AlphaFold pLDDT for various conditional generation experiments. A: Coulomb Bias on Lys with k=2000 with Length=20, B: Gaussian Bias on Asp with k=-2500 with Length=20, C: Constrain Sequences to Start with Met with Length=40, D: Net Charge Restrained at −1 with k = 2000 with Length=40. Error bars represent one standard deviation across the independently generated sequences.

## 5 Discussion

In this work we introduced ProtNHF, an NHF for controllable protein sequence generation. Our approach leverages deterministic, symplectic transport dynamics to learn a generative model of protein sequence embeddings, while enabling flexible conditioning at inference time through principled modifications of the learned Hamiltonian.

In the unconditional setting, the model produces high-quality protein sequences as assessed by ESM-2 pseudo-perplexity and AlphaFold2 structural confidence, demonstrating that Hamiltonian flows can serve as effective generative models for protein sequence space. These results establish that symplectic transport dynamics can faithfully model the complex distribution of natural protein sequences without requiring diffusion-based noise schedules or autoregressive factorization.

Crucially, we show that conditioning can be implemented without retraining by introducing smooth bias potentials into the Hamiltonian during sampling. This conditioning allows for protein sequences based on desired global sequence-level properties such as amino acid composition, and net charge. We empirically demonstrate three classes of conditioning:

- **Coulomb bias**, which can progressively promote or suppresses certain residues as the bias strength increases.
- **Gaussian bias**, which selectively enriches or depletes chosen residue types in a controlled, monotonic fashion.
- **Harmonic bias**, which enables positional residue control and global property targeting, including generation of sequences with prescribed net charge and greater secondary structure diversity.

Across all settings, conditioning strength produces predictable and continuous modulation of sequence statistics, indicating that energy shaping induces stable and controllable deformations of the learned transport map. Importantly, these modifications require no retraining and preserve the structural integrity of generated sequences.

Conceptually, our results demonstrate that conditional protein generation can be achieved by controlled perturbations of a learned Hamiltonian system. Rather than retraining a conditional model, our method modifies the transport dynamics itself, producing a new generative distribution via symplectic deformation. This distinguishes our approach from diffusion-based guidance, classifier-free guidance, and fine-tuning strategies, and highlights a unique advantage of Hamiltonian generative modeling: conditioning through energy shaping. Manipulation of residue-level properties constitutes protein engineering and as such these generative models could have important applications in the design of new artificial proteins and functional biomolecules. More broadly, this work establishes a connection between generative modeling and biased dynamical systems, showing that ideas inspired by molecular dynamics, such as bias potentials and energy shaping, can be repurposed for structured generative control in biological sequence space. This opens avenues for incorporating richer physics-inspired constraints, including structural priors, electrostatics-aware objectives, and functional site targeting, directly into generative dynamics.

In summary, we demonstrate that NHFs provide a mathematically principled and practically effective foundation for controllable protein generation, combining high-quality unconditional modeling with flexible, inference-time conditioning through energy-shaped transport.

## 6 Conclusions

Controllable protein sequence generation remains a central challenge in computational protein design. Most existing generative approaches achieve conditioning through retraining, classifier guidance, or architectural modification, limiting flexibility and increasing computational cost. Here we introduce ProtNHF, a neural Hamiltonian flow framework trained on the UniProtKB databased that enables high-quality protein sequence generation with conditioning applied exclusively at inference time. Specifically, it allows for conditional generation of protein sequences based on global sequence-level properties such as amino acid composition, and net charge. Manipulation of such properties constitutes protein engineering and is becoming increasingly important for the design of new artificial proteins and functional biomolecules.

Our model learns a symplectic transport map from a latent Gaussian distribution to protein sequence embeddings using deterministic leapfrog dynamics. In the unconditional setting, generated sequences achieve competitive quality as measured by ESM-2 pseudo-perplexity and AlphaFold2 pLDDT confidence scores, demonstrating that Hamiltonian flows can faithfully model protein sequence distributions.

We show that conditioning can be implemented without retraining by augmenting the learned Hamiltonian with smooth bias potentials during sampling. This energy shaping induces a controlled deformation of the learned transport map, yielding a new generative distribution while preserving deterministic, volume-preserving dynamics. We empirically validate three classes of conditioning: (i) Coulomb bias to suppress positively charged residues such as lysine, (ii) Gaussian bias to enrich or deplete selected residue types in a continuous and monotonic manner, and (iii) harmonic bias to enforce positional residue constraints and target global sequence properties such as net charge. In all cases, increasing bias strength results in predictable modulation of sequence statistics while maintaining structural plausibility.

Our results establish neural Hamiltonian flows as a principled and flexible framework for controllable protein generation. By enabling inference-time conditioning through dynamical perturbation rather than retraining, this approach provides a new connection between generative modeling and biased dynamical systems, opening avenues for physics-inspired control mechanisms in biological sequence design. Future work may explore theoretical guarantees on stability under strong biasing, the intergation of larger transformer architectures for more expressive modeling, and carrying out the flow in ensembles other than NVE. Extending this framework to structure generation or joint sequence-structure modeling represents another promising direction.

## 7 Acknowledgments

This research used resources of the Oak Ridge Leadership Computing Facility, which is a DOE Office of Science User Facility supported under Contract DE-AC05-00OR22725.

## 9 Data Availability

The code for ProtNHF is made available at https://github.com/bharath-raghavan/ProtNHF under the BSD-3 License. Trained model weights are available on Hugging Face at https://huggingface.co/bharathraghavan/ProtNHF, also under the BSD-3 License. Data related to all experiments discussed in this paper are available at https://doi.org/10.13139/OLCF/3019470.

